# Data-modeling the interplay between single cell shape, single cell protein expression, and tissue state

**DOI:** 10.1101/2024.05.29.595857

**Authors:** Yuval Tamir, Yuval Bussi, Claudia Owczarek, Luciana Luque, Giuseppe Torrisi, Leor Ariel Rose, Orit Kliper-Gross, Chris Sander, Linus Schumacher, Maddy Parsons, Leeat Keren, Assaf Zaritsky

## Abstract

Changes in cell shape are fundamentally involved in signaling, intracellular organization, function, and intercellular interactions within tissues, in health and disease. Investigating the interplay between cell shape and protein expression was limited, until recently, by the number of proteins that can be imaged simultaneously or by population averaging. We combined spatial multiplexed single cell imaging and machine learning to systematically investigate the intricate relationships between cell shape and protein expression in the context of heterogeneous human cells in their native state in human tissue samples in situ. Our analysis established a universal bi-directional link between the cell’s shape and its protein expression across different cell types, diseases, and disease states in human tissues, enabling new applications. Machine learning interpretability showed that the contribution of shape features to a prediction can potentially infer new protein functions. Unbiased screening of the links between all pairs consisting of one protein and one cell type identified a subpopulation of large p53-positive tumor cells across two cancers. Ultimately, inclusion of single cell shape properties enhanced Graph Neural Network disease state prediction. Our results open the door to unraveling the intricate connections between protein expression at the single cell level, cell shape, tissue organization, and tissue state in a physiological context.

## Introduction

Since the early days of Rudolf Virchow, and for over 200 years, pathologists have been using microscopes to observe the tissue’s architectural pattern, cellular rearrangement, cell and nuclear shape and intracellular organization to diagnose and make treatment decisions. Throughout evolution, diverse cell shape and intracellular organization, broadly termed *cell morphology*, enable diverse cellular functions (Mogilner & Keren, 2009; Mostafa, 2022). Single cell morphology, even when considering a single cell type, encodes information related to the gene expression patterns (Haghighi et al., 2022), signaling (Bakal et al., 2007a), functional states (Clark & Paluch, 2011), dynamics (K. Keren et al., 2008), and even metastatic potential (P.-H. Wu et al., 2020). Accordingly, single cell morphology, and specifically cell shape (Pincus & Theriot, 2007; Viana et al., 2023), are quantitative readouts extensively used for applications of high-content cell imaging such as systematic morphological profiling (Bakal et al., 2007a; Nassiri & McCall, 2018; Neumann et al., 2010) and drug screening (Caicedo et al., 2017; Chandrasekaran et al., 2021).

Recent spatial single cell imaging technologies, such as multiplexed ion beam imaging by time-of-flight (MIBI-TOF) (L. Keren et al., 2019) or Co-Detection-by-inDEXing (CODEX) (Black et al., 2021), enable multiplexed imaging of dozens of proteins at single cell resolution within tissue sections. These technologies enable to resolve different cell types and their spatial organization in their native context within the tissue, and to ask how the tissue’s spatial organization influences function and/or pathology. For example, several recent studies demonstrated that the cellular local neighborhood context is linked to the single cell’s protein expression profile (Fischer et al., 2023) and even to the disease state (Z. Wu et al., 2022). These spatial single cell imaging technologies enable to systematically study the fundamental relationships between cell shape and protein expression in physiological context in situ.

Here, we established that single cell shape and protein expression are inter-linked in disease-related tissue sections. Specifically, we showed that cell shape explains some of the within-cell type variation in protein expression and vice-versa in a human cohort of triple-negative breast cancer (TNBC) tissue sections. This link between single cell shape and protein expression was further confirmed for independent patient cohorts of tuberculosis (TB), colorectal cancer (CRC), melanoma, and head and neck cancer (HNC). By screening the links between cell shape and protein expression in all pairs consisting of one protein and one cell type in the TNBC cohort, we identified enrichment of the protein p53 in larger tumor cells.

This relation between p53 and tumor cell size was confirmed in another TNBC cohort that was acquired in a different hospital with a different technology. Finally, we showed that inclusion of single cell shape properties improves disease state prediction by graph neural networks that rely on the cells’ protein expression and their spatial organization in the TNBC and HNC cohorts. Altogether, our results indicate that cell shape contains information regarding the within cell type protein expression variations, and suggest that this information can be used to identify new cell subtypes and cell states, and to enhance the prediction of disease state in heterogeneous tissue sections.

## Results

### Cell shape heterogeneity in triple negative breast cancer (TNBC)

We reanalyzed data from a human TNBC cohort, previously collected by MIBI-TOF, consisting of 40 tissue sections from different patients depicting 36 protein channels at subcellular resolution (L. Keren et al., 2018). Using the single cell segmentation masks (Van Valen et al., 2016), we extracted 12 standard shape features, and expert-validated annotations categorized every cell to one of 16 different cell types (Fig. 1A). Visual assessment showed heterogeneity in the cell shapes, even within the same cell type (Fig. 1A). Systematically associating cell shape to cell type showed that tumor cells were the largest and the most heterogeneous cell type, Macrophages had the lowest convex-hull to area ratio, and T-regulatory cells were the most homogeneous in terms of size (Appendix 1). However, the high cell shape variability between patients and within cell types induced only weak associations between cell type and cell shape (Appendix 1) indicating that cell shape does not contain strong discriminative information regarding the cell type, at least at the crude spatial resolution of 0.5 µm microns per pixel, that is standard in spatial single cell assays.

**Figure 1.**
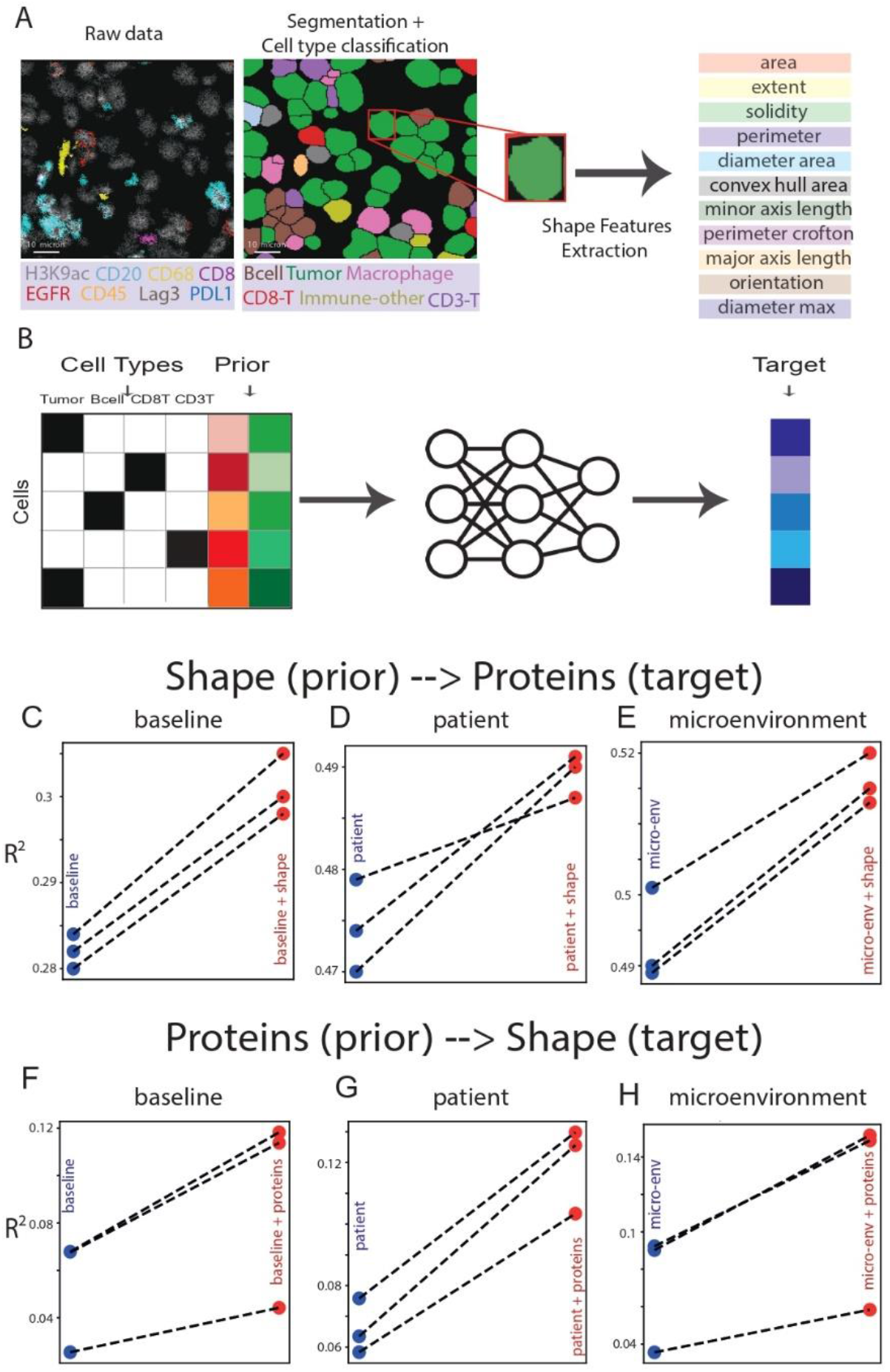
Single cell shape and protein expression are associated. **(A)** Preprocessing and shape features extraction. The raw multiplexed images were segmented to single cells, and each cell was categorized according to its cell type. Shape features were extracted from the single cell segmentation masks. **(B)** To isolate and quantify the influence of a prior (cell shape or protein expression) on the prediction of a single cell target (protein expression or cell shape, correspondingly) we compared the performance of a baseline model versus the prior-aware model. The schematic depicts the prior-aware model. Each row represents a cell, where the cell type is encoded via one hot encoding and the prior is concatenated to it. **(C-E)** Shape-aware models surpassed their corresponding baseline models in predicting single cell protein expression. Each matched data pair represents the 3-fold cross-validation R^2^ scores between the prediction and observed protein expression of the baseline (blue) and its matched shape-aware predictions (red). Statistical significance for the contribution of shape was calculated via paired t-tests comparing the R^2^ scores of patients between the baseline and the shape-aware model to test the null hypothesis that the matched differences between these models are distributed around 0. The null hypothesis was rejected with a p-value < 0.05 for all panels. **(C)** Baseline model using the cell type versus the corresponding shape-aware models. Per-patient analysis in Fig. S2A (p-value = 0.027). **(D)** Baseline model using the cell type and the patient (batch) versus the corresponding shape-aware models. Per-patient analysis in Fig. S2B (p-value = 0.034). **(E)** Baseline model using the cell type, the patient and the cell’s microenvironment composition versus the corresponding shape-aware models. Per-patient analysis in Fig. S2C (p-value = 0.013). **(F-H)** Protein expression-aware models surpassed their corresponding baseline models in predicting single cell shape. Each matched data pair represents the 3-fold cross-validation R^2^ scores between the prediction and observed cell shape features of the baseline (blue) and its matched protein expression-aware predictions (red). Statistical significance for the contribution of protein expression was calculated via paired t-tests comparing the R^2^ scores of patients between the baseline and the protein expression-aware model to test the null hypothesis that the matched differences between these models are distributed around 0. The null hypothesis was rejected with a p-value < 0.05 for all panels. **(F)** Baseline model using the cell type versus the corresponding protein expression-aware models. Per-patient analysis in Fig. S2D (p-value < 0.001). **(G)** Baseline model using the cell type and the patient (batch) versus the corresponding protein expression-aware models. Per-patient analysis in Fig. S2E (p-value = 0.01). **(H)** Baseline model using the cell type, the patient, and the cell’s microenvironment composition versus the corresponding protein expression-aware models. Per-patient analysis in Fig. S2F (p-value < 0.001).

### Within cell type association of single cell shape and protein expression

Most of the MIBI-TOF’s 36 multiplexed protein antibodies are selected to determine the cell type. Thus, the main source of variability between the cells’ protein expression is attributed to the cell type. The residual variation in protein expression within a cell type is commonly attributed to cell cycle states (Gut et al., 2018), microenvironment (Fischer et al., 2023; E. Wu et al., 2023)), and a more refined cell state characterization (Alizadeh et al., 2020). Given the high variations in cell shapes within the same cell type we hypothesized that some of the variations in the normalized protein expression per area unit (from here on referred as “protein expression”) within the same cell type can be related to the cell shape, and vice versa -that some of the variation in cell shape can be related to the protein expression. We tested this hypothesis by using machine learning to predict the cell’s protein expression given its cell shape, and to predict the cell’s shape given its protein expression (Fig. 1B, in this context the cell shape or the protein expression are termed the “*prior*”). First, we assessed whether the inclusion of cell shape features (i.e. shape-aware model) contributed to the prediction of the protein expression beyond a baseline model defined by the cell type alone (Methods). Specifically, for each model, we calculated the coefficient of determination (R^2^), the standard metric for assessing the accuracy of a regression task, and the ΔR^2^ was used for model comparison (see methods). All shape-aware models surpassed the baseline model and specifically, a shape-aware fully connected deep neural network (DNN) model surpassed the baseline model in 36 out of the 40 patients’ tissue sections, underscoring that cell shape encodes some of the variations in protein expression beyond the information encoded by the cell type alone (Fig. 1C, per-patient results in Fig. S2A). The DNN model also surpassed shape-aware linear models (Fig. S1). To exclude the possibility that this contribution of cell shape to the prediction of protein expression was confounded by inter-patient variability, we validated that cell shape contributes to a model conditioned on both the cell type and the patient (i.e., batch, see methods). The shape-aware model surpassed the model without shape information in 35 out of the 40 patients (Fig. 1D, Fig. S2B). To exclude the possibility that the residual protein expression explained by cell shape was confounded by the cell microenvironment, we validated that inclusion of cell shape surpassed a model that was trained with the cell type, patient and the cell microenvironment composition in 35 out of the 40 patients. The microenvironment composition was defined for every cell using a radius of 70 µm as ((Fischer et al., 2023), Methods) (Fig. 1E, Fig. S2C). In all cases cell shape showed a modest but significant contribution to the prediction of protein expression to the prediction of protein expression, with an average ΔR^2^ of ∼2%. This magnitude of improvement is in agreement with a previous study that explored the contribution of the cellular microenvironment to the prediction of single cell protein expression (Fischer et al., 2023), aligning with the understanding that the most dominant variability factor in the protein expression is the cell type.

Having established that cell shape contributes to the prediction of protein expression beyond cell type, we investigated the opposite direction to determine whether protein expression provides information beyond cell type for cell shape prediction. Similarly to the previous analysis, we trained a fully-connected neural network model to predict the 12 cell shape features from the cell’s protein expression and compared its performance to a baseline model defined by cell type alone. The protein expression-aware model surpassed the baseline cell shape model with an average ΔR^2^ of ∼5% (Fig. 1F, Fig. S2D), and this contribution of protein expression to cell shape prediction was not confounded by patient identity (Fig. 1G, Fig. S2E) nor the cellular microenvironment composition (Fig. 1H, Fig. S2F). These results indicate that the protein expression encodes information about cell shape beyond what is captured by cell type.

Our analysis of the intra-cell type variations suggests an association between cell shape and protein expression in TNBC tumors. To assess the generalizability of these findings, we repeated the same analysis on three additional datasets: a public tuberculosis MIBI-TOF dataset (McCaffrey et al., 2022), a public CODEX colorectal cancer (CRC) dataset (Schürch et al., 2020) and a Melanoma MIBI-TOF dataset (Methods). Across all three datasets, shape- and protein-aware models surpassed their corresponding baseline models (Fig. S4). These results suggest an association between cell shape and protein expression that generalizes across multiple human diseases.

### Shape features importance cluster according to proteins function

Cell shape was found to contribute to the protein expression prediction, but it was not clear which of the shape features were the ones most influential in improving the predictions. To explain the predictions of the shape-aware model we applied the SHapley Additive exPlanations (SHAP) interpretability method (Lundberg & Lee, 2017a) that assigns each feature an importance value for each prediction of our shape-aware baseline model. For the purpose of evaluating shape, we ignored the SHAP values attributed to the cell type feature, focusing on the SHAP values attributed to the 12 cell shape features. We aggregated the SHAP explanations for each protein to find that size-related features, such as perimeter and area, had the highest overall importance, and shape-related features, such as orientation, solidity, and eccentricity were slightly less important for the model’s predictions across all protein targets (Fig. S3A). We performed hierarchical clustering of the proteins according to their corresponding shape’s 12-dimensional SHAP values. This analysis identified clusters of proteins whose predictions were similarly influenced by the cell shape (Fig. S3B). One cluster included Keratin-6, Beta-catenin, Keratin 17, and pan Keratin, proteins that are known to be overexpressed in many cancers including TNBC and serve as potential diagnostic and prognostic markers (Baraks et al., 2022; Maeda et al., 2016; Menz et al., 2023; Weeks et al., 2021). A second cluster comprised of H3K9ac, dsDNA, H3K27me3, Ki67, and Phospho-S6, which includes nuclear cell status markers for proliferation (Ki67, Sun & Kaufman, 2018), and for histone modification markers (H3K27me3 and H3K9ac, Igolkina et al., 2019). Zooming in to the SHAP values of the individual proteins showed striking within-cluster similarities and between-cluster differences (Fig. S3C). These results suggest that cell shape importance is associated with the protein function and thus may be used to infer new potential protein functions.

### Tumor cell shape is associated with p53 expression

We next used the same approach to measure more specifically the contribution of cell shape to the prediction of protein expression in the context of the cell type. For each pair of (cell type, protein) we measured the residual variability in the protein’s expression within the cell type that is explained by the cell shape (Fig. 2A). Namely, averaged across patients for every pair of (cell type, protein), we constructed the ΔR^2^ matrix, where for each bin (*i,j*) we calculated the contribution of the shape to the prediction of the expression of the specific protein *i* in the context of cell type *j* . This ΔR^2^ matrix encoded 16 (cell types) x 36 (proteins) = 576 such relations, enabling us to screen for shape-dependent proteins in the context of their cell type (Fig. 2B). Expectedly, cell shape contributed to the prediction of the expression levels of dsDNA and histones (HEK9ac, HEK2me3), presumably reflecting coordinated changes in both cell shape and histone content during the cell cycle (Nagao et al., 2020; Roukos et al., 2015). Next, we examined the contribution of the cell shape to the expression of p53 in tumor cells. p53 mutations are frequently observed in many types of cancers, including TNBC, and the protein plays a role in regulating cell cycle arrest, apoptosis, and senescence in response to various cellular stresses (Hafner et al., 2019). Heterogeneity in p53 expression within tumor cell subpopulations (Yang et al., 2013) was speculated to influence their behavior and response to therapy (Li et al., 2015). In the context of TNBC, p53 mutations are highly prevalent, with mutation rates reaching up to 80% (Cancer Genome Atlas Network, 2012), often leading to the accumulation of nonfunctional p53, and contributing to the aggressive nature and poor prognosis of TNBC (Bianchini et al., 2016).

**Figure 2.**
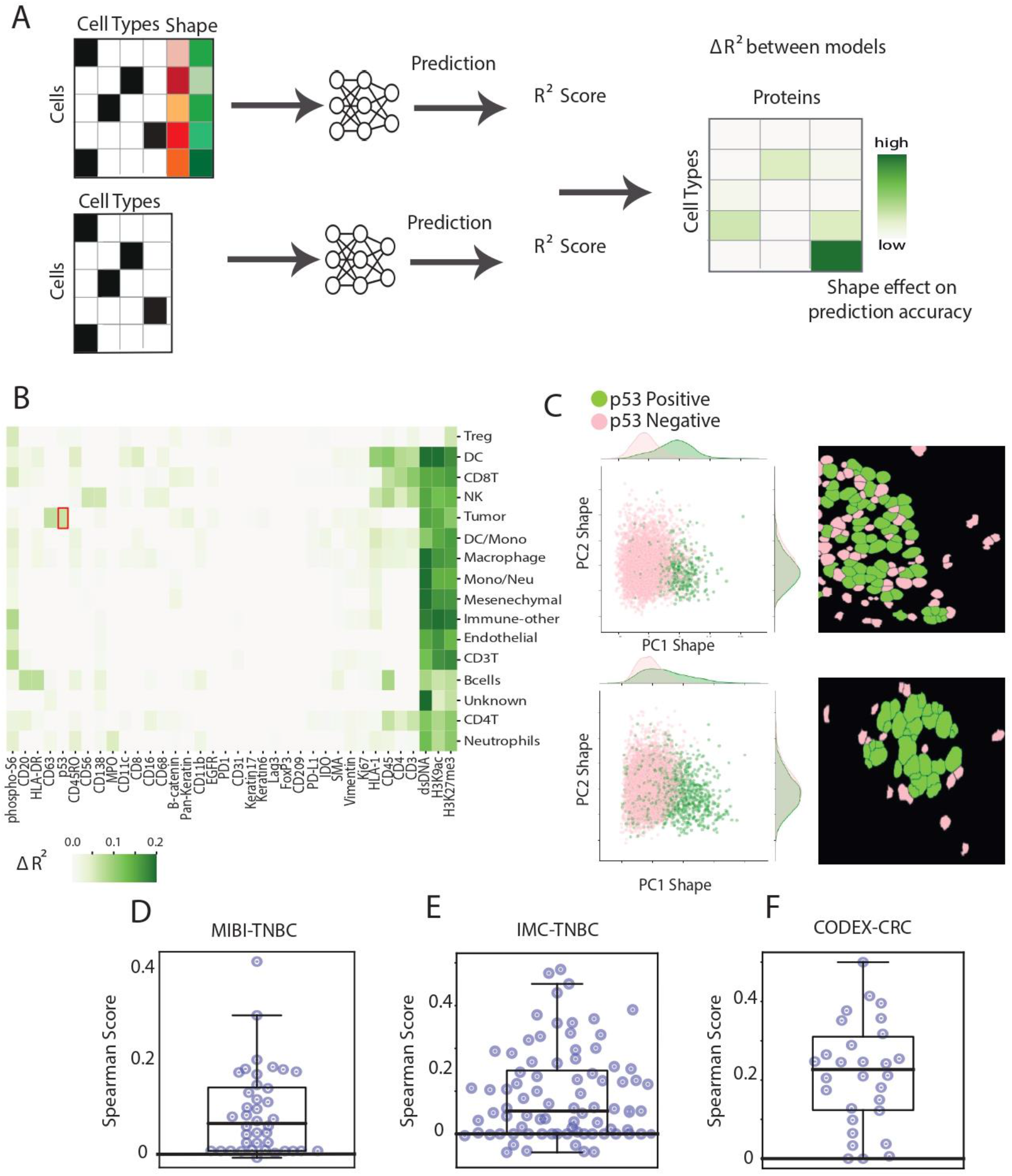
Screening for shape-dependent protein expression. **(A)** For each pair of cell type and protein expression we calculated the change in the R^2^ values between the observed and the predictions of the shape-aware model in respect to the baseline. These pairs define the ΔR^2^ matrix that measures and visualizes the contribution of cell shape to the prediction of the protein expression (x-axis) in the context of a cell type (y-axis). **(B)** Visualization of the ΔR^2^ matrix computed by subtracting the baseline model’s R^2^ scores from the shape-aware model R^2^ scores. For visual clarity only positive values (508/576) are displayed. The x-axis represents the quantified proteins, and the y-axis represents the annotated cell types. The red box emphasizes the shape-dependent improvement in tumor cells’ p53 expression prediction. **(C)** Representative examples of two patients (#28 top, #16 bottom) demonstrating the association between p53 positive/negative status (determined according to the p53 expression, see Methods) and cell shape. Left: Scatter plots of tumor cells’ 2D shape-space principal component analysis (PCA) projection, with cells colored according to their p53 positive (green) or negative (pink) status. A t-test rejected the null hypothesis that the PC1 values are distributed around the same mean for both p53-positive and p53-negative cells with a p-value < 0.0001. Statistical significance was recorded in 16/20 of patients that had a sufficient cell count (>100) of p53-positive and p53-negative tumor cells (Fig. S5). Right: Representative segmented tumor cells from the corresponding patients show apparent qualitative shape difference between p53-positive (green) and p53-negative (pink) tumor cells. **(D-F)** Spearman correlation scores between the cell’s p53 expression and its area across different patients (each represented by a data point), derived from a MIBI-TOF TNBC with N = 40 patients (**D**), an IMC TNBC with N = 58 patients (**E**), and a CODEX CRC with N = 35 patients (**F**) cohorts. All panels exhibited a similar pattern, with data points above 0, indicating a positive correlation between p53 expression and cell size. For all panels, one sample t-test p-values < 0.0001, suggesting strong evidence against the null hypothesis of no correlation. For visualization purposes, one outlier patient was discarded from the IMC TNBC dataset.

Here, we revealed an association linking high expression of p53 in tumor cells and cell shape (Fig. 2B). This association between p53 and tumor cell shape was visually and statistically evident in some patients as demonstrated by a separation in the two-dimensional PCA-reduced shape space when comparing p53-positive and p53-negative (as defined in (L. Keren et al., 2018b)) cells (Fig. 2C, Fig. S5). Visual observation (Fig. 2C, right) and quantification (Fig. 2D) showed that p53-positive cells were larger than p53-negative cells suggesting the existence of a p53-expression-dependent subpopulation with distinct size signatures. We confirmed this association in an independent cohort of TNBC patients from a different clinic, acquired by a different lab using a different technology (Imaging Mass Cytometry, IMC) (Methods) (Fig. 2E). Finally, we revealed that this size-associated p53 expression of tumor cells was also apparent in a cohort of colorectal cancer patients (Schürch et al., 2020) (Fig. 2F). Overall, we suggest that p53 expression in tumor cells is associated with increased cell size across a variety of cancers.

### Cell shape contributes to Graph Neural Network-based clinical prediction

Our results indicate that cell shape features contribute to the quantitative description of protein expression within TNBC tissues, but can it contribute for better characterization of disease state? To investigate the potential contribution of shape features to the prediction of clinical outcomes, we defined a graph classification problem (see Methods). Specifically, we constructed an undirected network graph based on the image data, as outlined in (Z. Wu et al., 2022), where each cell represented a node, with its protein expression as the node’s attributes. The network was then fed into a Graph Convolutional Network (GCN), which is a specialized type of neural network designed to work directly with graph structures (Kipf & Welling, 2016) (Fig. 3A). The GCN operates by propagating information between connected nodes in the graph, effectively enabling each node to “gather” information from its neighboring nodes. This process allows the network to learn both local and global patterns in the data. Each layer in the GCN captures increasing levels of neighborhood information, enabling nodes to “communicate” with nodes farther away in the graph. Previous studies showed that this approach could predict clinical outcomes, with cohorts in the order of dozens of patients (e.g., (Z. Wu et al., 2022).

**Figure 3.**
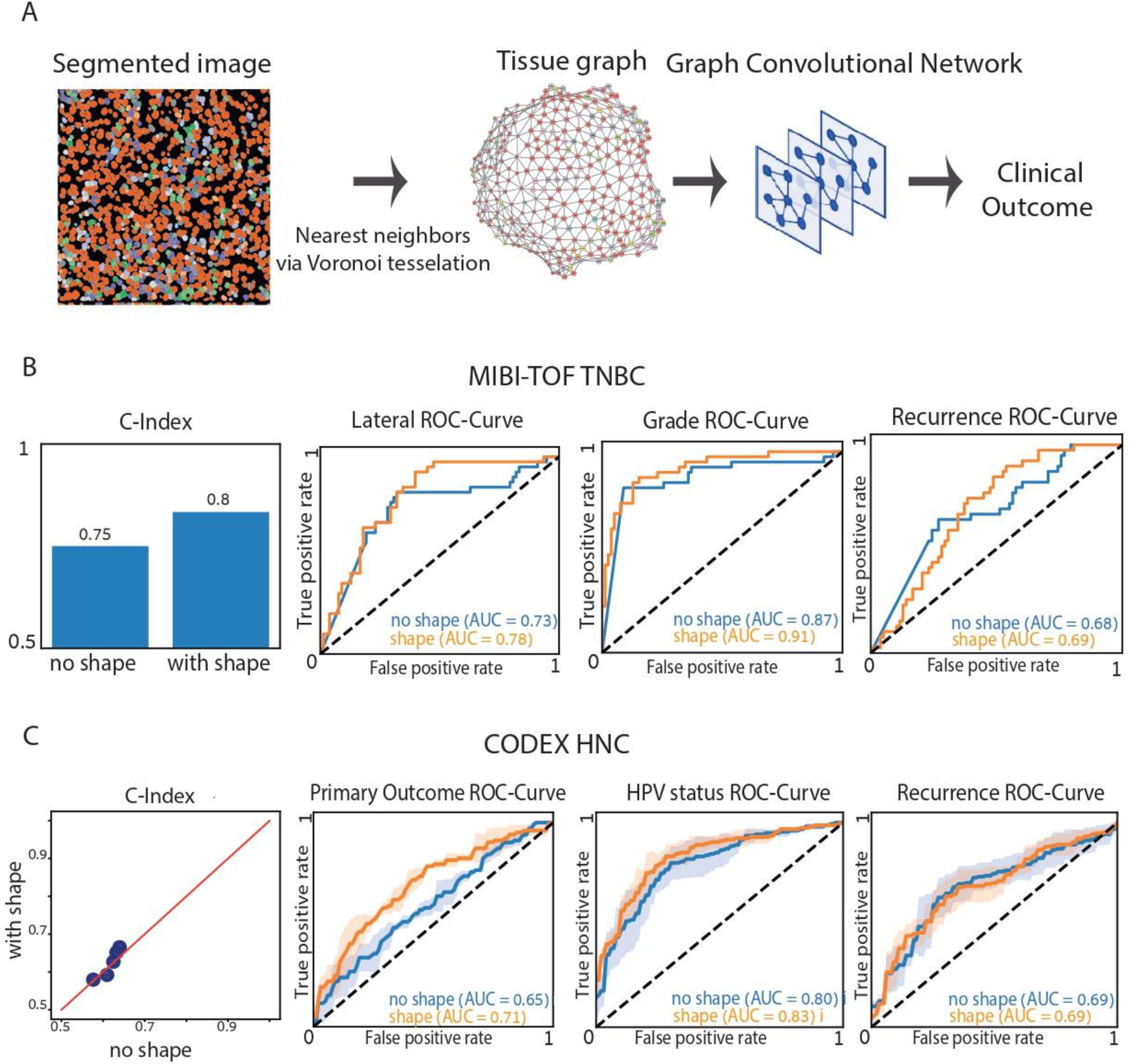
Cell shape contribution to the prediction of the patients’ clinical outcome. (**A**) Computational pipeline (left-to-right): (i) input: cell segmentation and cell type classification, (ii) cell adjacency network (aka “graph”): cells are nodes and Delaunay neighbors define edges, node features are defined as the cells protein expression with or without shape features, (iii) Graph Convolutional Network maps the graph to learn the outcome clinical label. (**B-C**) Cell shape contributes to prediction of clinical outcome. The ROC curves compare the performance of shape-aware models (orange) and shape-independent models (blue). Shape-aware GCNs led to an improvement in most clinical prediction tasks. **(B)** TNBC cohort (N = 40 patients) clinical outcome (left-to-right): C-index -lifespan post-diagnosis (survival ranking metric), lateral - edge of the tissue (binary measure), Grade - tumor aggressiveness level (multiclass), Recurrence - returning cancer (binary measure). For Grade, the micro-average ROC (weighted average) is presented due to the highly imbalanced classes. Due to the limited number of patients, the models were evaluated using a leave-one-out strategy, where each sample was iteratively held out as a test set while the remaining samples were used for training. **(C)** Head and neck (HNC) (N = 139 patients) clinical outcome (left-to-right): C-index - lifespan post-diagnosis (survival ranking metric), Primary outcome - squamous cell carcinomas (HNSCC) (positive or negative), HPV status - presence of human papillomavirus (binary), Recurrence - returning cancer (binary). The models were evaluated using 5-fold cross-validation. In the C-index scatter plot (left), each data point represents one of the 5-fold cross-validation scores of a shape-aware (y-axis) and baseline (x-axis) predictions. Data points above the diagonal y = x indicate that the corresponding shape-aware model induces more accurate predictions. ROC curves (right) show the mean (solid line) and the standard deviation (shade) of the shape-aware (orange) and baseline (blue) models’ performance.

We demonstrated that GCNs could predict the disease state from the tissue sections’ single cell networks for three cohorts (MIBI-TOF TNBC, IMC TNBC and CODEX HNC) and for multiple clinical readouts (Fig. 3B-C, Fig. S6). To evaluate the contribution of the cell shape to the clinical outcome prediction, we compared the performance of two models: 1) a baseline GCN model as described above, and 2) the baseline model with the inclusion of single cell shape as a conditioning factor. The shape-aware GCN models surpassed their corresponding baseline models in predicting a total of 8 different clinical readouts from these three datasets (Fig. 3B-C, Fig. S6). These results indicate that single cell shape encodes complementary discriminative information regarding the disease state, beyond the protein expression and the cells’ spatial composition.

## Discussion

Using disease-related human tissue sections we systematically investigated the bi-directional link between the cell’s shape and its protein expression in the context of multiple cell types, diseases and disease states. Cell shape is precisely regulated by a combination of intracellular and extracellular cues, can control cell signaling, is indicative of the cell’s physiological state, and its dysregulation can be linked to disease progression stage (Bakal et al., 2007b; Cachoux et al., 2023; Dent et al., 2024; Driscoll et al., 2019; Goudarzi et al., 2017; Gut et al., 2018; Haftbaradaran Esfahani & Knöll, 2020; Kai et al., 2016; K. Keren et al., 2008; Lamouille et al., 2014; Marshall, 2020; McBeath et al., 2004; Meyers et al., 2006; Mills et al., 1998; Qu et al., 2019; Rohban et al., 2017; Sailem & Bakal, 2017; Segal et al., 2022; Tee et al., 2011; Viana et al., 2023; Weems et al., 2023; P.-H. Wu et al., 2020; Yanagida et al., 2022; Yin et al., 2013). Accordingly, cell shape is a key readout for pathological diagnosis and treatment decisions, and one of the most studied readouts in the broad domain of cell biology. Until recently, the ability to systematically study the relationships between cell shape and gene expression was restricted to population averages of cell line models (Haghighi et al., 2022). The recent emergence of spatial multiplexed single cell imaging technologies enabled us to quantitatively study the intricate relationships between the shape of single cells and their protein expression within a more physiologically relevant context of heterogeneous cell populations in their native state of intact tissue samples in situ. Specifically, using machine learning performance as our metric we established that some of the within-cell-type protein expression variability can be explained by the cell’s shape and some of the within-cell-type cell shape variability can be explained by the cell’s protein expression. These associations between cell shape and protein expression seem to be universal across cohorts of four different diseases (TNBC, TB, CRC, NHC) acquired using two technologies (MIBI-TOF, CODEX).

We demonstrate multiple applications where integration of cell shape can contribute to downstream analyses of spatial multiplexed single cell imaging data. First, identification of new cell subtypes and cell states. We showed that the contribution of cell shape to protein expression prediction can unbiasedly screen all matched cell-type and protein pairs. In the context of TNBC, such a screen identified (in one cohort) and then confirmed (in another cohort) an enrichment of p53 expression in larger tumor cells, a subpopulation that was also identified in a CRC cohort. Second, generation of new insight regarding protein functionalities. We showed that the interpretation of the most prominent shape features toward the prediction of a protein expression uncovered shape signatures that were clustered according to the protein functions, highlighting the potential of using shape-based representations to infer new protein functions. Third, enhancement of clinical outcome prediction. We showed that inclusion of shape features improved Graph Convolutional Network-based predictions across three cohorts with multiple clinical readouts per cohort.

The terms cell “shape” and “morphology” are sometimes used interchangeably, but have a distinct meaning. Cell morphology includes cell shape, but encompasses a broader set of measurements that encode a richer representation of the cell state. Morphological readouts can include information regarding the intracellular organization and the intercellular context beyond that encoded in cell shape, for example metabolic and proteomic components by implicitly extracting this information from the raw image texture. In the context of spatial multiplexed technologies, cell morphology has shown to assist in mapping 7-to-40-plex CODEX at single-cell resolution (E. Wu et al., 2023), in cell type classification (Amitay et al., 2023). The downside of cell morphology is that it is more difficult to biologically interpret and thus it is harder to extract specific biologically meaningful insights. Cell shape is widely studied and can be more easily interpreted (e.g., larger p53-positive tumor cells), validated and linked to specific molecular players (e.g., Jones et al., 2023) and thus was at the focus of our study.

The prediction performance gain attributed to cell shape, was consistent across all datasets and applications, but moderate in terms of its magnitude. A similar performance gain was recently reported for the contribution of the cell’s microenvironment to the prediction of single cell protein expression beyond the cell type (Fischer et al., 2023b). One reason for this modest effect is that the cell type is the most dominant factor in explaining the variability in the protein expression, especially in current panels that are limited to a few dozens of proteins, where most of them are used to infer the cell type. Technological advancements are expected to increase the number of proteins enabling larger numbers of “cell state” markers that we expect to have higher associations with the cell shape (Y. Liu et al., 2023; Scheuermann et al., 2024). Another reason for the moderate contribution of the cell shape contribution is the low spatial resolution. Physical pixel sizes of 0.4-1 µm, which are a standard in spatial multiplexed technologies, are limiting (even perfect) cell segmentation to a rough outline. This is probably the main reason behind our failure to predict cell type from cell shape. On the other hand, we argue that the ability to consistently measure the contribution of cell shape to a variety of applications and datasets indicates a strong biological signal that goes beyond the noise levels introduced by the coarse segmentation. Indeed, high resolution and 3D cell shapes provide discriminative information regarding the cell’s microenvironment (Dent et al., 2024; Segal et al., 2022), molecular organization (Mazloom-Farsibaf et al., 2023; Viana et al., 2023) and treatment (De Vries et al., 2023). Also here, future technologies are expected to have improved resolution and maybe even 3D information leading to information-rich segmentations. These segmentations will enable the application of more powerful shape descriptors (e.g., Mazloom-Farsibaf et al., 2023; Viana et al., 2023) and enhanced discrimination between different cell types states.

## Methods

### Data

We analyzed four public and two new datasets (Supplementary Table 1).

#### Triple Negative Breast Cancer MIBI-TOF (MIBI-TNBC)

We reused an existing MIBI-TOF TNBC dataset that included 40 patients (L. Keren et al., 2018b). 36 proteins were used to identify 16 cell types across 178,366 cells, with physical pixel size of 0.5x0.5 µm2. We used the same cell types annotations (Tumor, Endothelial, Mesenchyme, Tregs, CD4 T cells, CD8 T cells, CD3 T cells, NK cells, B cells, Neutrophils, Macrophages, DC, DC/Mono, Mono/Neu, Immune other), and the same segmentation masks from (L. Keren et al., 2018b). Clinical readouts include: lifespan post-diagnosis, lateral (i.e., sample taken from the edge of the tissue, binary), grade (tumor aggressiveness level), and recurrence (binary).

#### Tuberculosis MIBI-TOF (MIBI-TB)

We reused an existing MIBI-TOF TB granulomas dataset that included 26 patients (McCaffrey et al., 2022). 38 proteins were used to identify 19 cell types across 56,402 cells, with physical pixel size of 0.5x0.5 µm^2^. We used the same cell type annotations (endothelial, CD16_CD14_Mono, CD8_T, CD4_T, imm_other, CD14_Mono, neutrophil, fibroblast, B_cell, CD68_Mac, CD206_Mac, CD11c_DC/Mono, CD11b/c_CD206_Mac/Mono, mast, CD163_Mac, Treg, and epithelial) and the same segmentation masks from (McCaffrey et al., 2022).

#### Head and neck cancer CODEX (CODEX-HNC)

We reused an existing CODEX head and neck cancer (HNC) dataset that included 139 patients (Z. Wu et al., 2022). 22 proteins were used to identify 16 cell types across 2,061,162 cells, with physical pixel size of 0.4x0.4 µm^2^. We used the same cell type annotations (Tumor (Podo+), Tumor (CD20+), CD4 T cell, Tumor (Ki67+), Stromal / Fibroblast, Tumor (CD21+), Naive immune cell, Macrophage, B cell, Tumor, APC, Lymph vessel, CD8 T cell, Vessel, Granulocyte, Tumor (CD15+)) and the same segmentation masks from (Z. Wu et al., 2022). Clinical readouts include: lifespan post-diagnosis, primary outcome (HNSCC positive or negative), HPV status (i.e., presence of human papillomavirus, binary), recurrence (binary).

#### Collateral cancer CODEX (CODEX-CRC)

We reused an existing CODEX Colorectal cancer (CRC) dataset that included 35 patients (Schürch et al., 2020). 56 proteins were used to identify 16 cell types across 218,372 cells, with physical pixel size of 0.4x0.4 µm^2^. We used the same cell type annotations (B cell, CD4T, CD8T, CD3T, Treg, DC, Fibroblast, Tumor, Macrophage, Neuron, Neutrophil, Plasma, Stroma, Endothelial, SMV, and Lymphatic) and the same segmentation masks from (Schürch et al., 2020).

#### Melanoma MIBI-TOF (MIBI-Melanoma)

We analyzed an unpublished MIBI-TOF dataset of melanoma that included 69 patients (multiple ROIs per patient). 36 proteins were used to identify 27 cell types and subtypes across 1,701,560 cells, with a physical pixel size of 0.5x0.5 µm^2^. The cell types identified in this dataset include DC sign Mac, blood vessels, Unidentified, Collagen_sma, B cell, CD4 APC, CD4 T cell, CD20_neg_B_cells, SMA, CD8 T cell, Mac, Collagen, Memory_CD4_T_Cells, CD206_Mac, Neutrophil, NK cell, Mono_CD14_DR, CD11_CD11c_DCsign_DCs, CD68_Mac, Hevs, CD4 Treg, CD14_CD11c_DCs, DCs, Follicular_Germinal_B_Cell, Tfh, Immune, and Tumor. Segmentation masks were generated using DeepCell2 (Bannon et al., 2021). Cell types were classified with CellSighter (Amitay et al., 2023).

#### Triple Negative Breast Cancer IMC (IMC-TNBC)

We analyzed an unpublished Imaging Mass Cytometry (IMC) dataset of Triple-Negative Breast Cancer (TNBC) from 58 patients (multiple ROIs per patient). 4-μm-thick sections from FFPE blocks were stained with a panel of 35 antibodies, targeting the main cell populations including immune, stromal, and tumour/epithelial cells (Supplementary Table 1). Slides were incubated for 1 h at 60°C, dewaxed, rehydrated, and then subjected to antigen retrieval using citrate buffer (pH 9, 96^0^C, 3 minutes). Slides were incubated in 10% BSA (Sigma), 0.1% Tween (Sigma), 5 mg/ml IgG (Kiovig, Shire) and 5% FcR in Superblock blocking buffer (Thermo Fisher) at room temperature for 2 h. Combined antibodies were mixed in blocking solution and incubated overnight at 4°C. Slides were washed twice in PBS-Tween and incubated for 30 min with the 1.25 mM (in PBS) of the DNA intercalator Cell-ID™Intercalator-Ir (Standard BioTools) (isotopes 191Ir and 193Ir). Slides were then washed in PBS and MilliQ water followed by air-drying. Samples were imaged using the Hyperion Imaging System (Standard BioTools) imaging module to obtain a light-contrast high-resolution image of each stained section. These images were used to select the region of interest (ROI) in each slide for ablation at a 1 μm/pixel resolution and 200 Hz.

The dataset contained 3,738,846 cells, with a physical pixel size of 1x1 µm^2^. The cell types identified in this dataset include Exhausted T-cell, Immune/Cancer, Th1 T helper, CAF, NK cell (FOXP3), Monocyte (Tbet), GB Active T cell, Macrophage, T-cell, B7H4+ Cancer (CD11b), Unassigned, Endothelial, Monocytes, Memory T cell, Monocyte (HLA), B/T/NK cell, CD44+ MAC, B7H4+ Cancer, Vim+ Cancer / NK, Mem CD8 T cell, Cancer stem cell, Antigen presenting cell, Reg T-cell, Cytotoxic T cell, E-Cad cancer, FOXP3+ Cancer, NK cells / Cancer, T/B cells, PanCK Cancer, B-cell, Alfa-SMA+ Monocytes, and T-cell/Cancer. Data pre-processing was done using steinbock (Windhager et al., 2023), segmentation masks were generated using DeepCell2 (Bannon et al., 2021), and cell types were clustered and defined using Pixie (C. C. Liu et al., 2023). Clinical readouts in the data: treatment response (binary).

## Shape features

We used python’s skimage library (region props function) to extract 12 typical shape features for each cell’s segmentation mask (Carpenter et al., 2006; C. C. Liu et al., 2023). For full details see Appendix Table 1.

### Machine learning models

For all our deep learning models, i.e. Deep Neural Network (DNN) and Graph Convolutional Networks (GCN) we used python’s pytorch and pytorch geometric libraries. For linear models we used scikit-learn linear models (LinearRegression, Ridge and Lasso). 3-fold cross validation was performed with the DNN, and 5-fold cross validation was performed with the GCN. In each fold, a different subset of the cells was chosen randomly for the test.

#### Baseline model

A three-layer DNN *f*_*baseline*_ is trained to predict a cell’s target *y* ∈ ℝ^*m*^ from its one-hot encoded cell-type vector *x*_*cell*_ ∈{0,1}^*n*^.*n* represents the number of cell types, and signifies the dimensions of the target. The learned model parameters are noted as Θ. The model function is given by *f*_*baseline*_(*x*_*cell*_ ; Θ) . Note that the baseline model predicts a continuous target (protein expression or shape, Fig.1) from a discrete input (cell type) and thus loses the within-cell type heterogeneity. Thus, we expect the prediction to be close to the mean of the cell type class’s target.

### Prior-aware model

Prior-aware models enhances *f*_*baseline*_ by incorporating prior features (shape or protein expression). Denoted by *x*_*prior*_ ∈ ℝ^*p*^, the prior is concatenated to the one-hot encoded cell-type vector *x*_*cell*_ ∈{0,1}^*n*^. Here, *p* represents the prior features dimensions (*p* = 12 for cell shape, e.g., *p* = 36 for the MIBI-TNBC data). The model function is given by *f*_*baseline + prior*_ (*x*_*cell*_ | | *x*_*prior*_ Θ) = *y*.

### Patient-aware model

Patient-aware (and called ‘batch’) model enhances *f*_*baseline*_ by incorporating patient ID identifier *x*_*patient*_ ∈{0,1}^*b*^ alongside the one-hot encoded cell-type vector *x*_*cell*_ ∈{0,1}^*n*^, aiming to capture patient-specific variation in the target (protein expression or shape). Here, *b* represents the number of patients. The model function is given by:

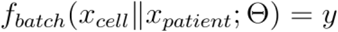

### Patient- and prior-aware model

Combines *x*_*cell*_ ∈ {0,1}^*n*^, *x*_*patient*_ ∈ {0,1}^*b*^ and *x*_*prior*_ ∈ ℝ^*p*^. The model function is given by: *f*_*batch + prior*_ (*x*_*cell*_ | | *x*_*patient*_; | |*x*_*prior*_; Θ) = *y*.

### Microenvironment-aware model

Following the work of (Fischer et al., 2023b), the microenvironment-aware models input is composed of *x*_*cell*_ ∈ {0,1}^*n*^, *x*_*patient*_ ∈ {0,1}^*b*^, and a binary microenvironment representation *x*_*microenvironment*_ ∈ {0,1}^*l*^, indicating the presence of each cell type within a fixed radius of 70 µm around the target cell. is the number of cell types, and the binary digit indicates whether the specific cell type is within the specified radius, not taking into account the number of cells. This model integrates spatial context to predict the target. The microenvironment-aware model function is given by:

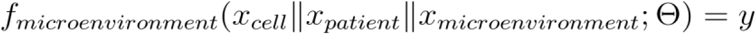

### Microenvironment- and prior-aware model

Combines *x*_*cell*_ ∈ {0,1}^*n*^, *x*_*patient*_ ∈ {0,1}^*b*^, *x*_*prior*_ ∈ ℝ^*p*^, and *x*_*microenvironment*_ ∈ {0,1}^*l*^. The model function is given by:

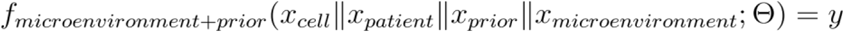

Note that in all models, the prior is either cell shape, or cell protein expression, and the target is either protein expression or cell shape, accordingly, depending on the model configuration (Fig. 1)

### Graph Convolutional Network

In this model we perform a whole graph classification task to predict the clinical outcome, using a 3 layers graph convolutional network (GCN). For this purpose, each tissue section was transformed to an undirected graph, where cells defined the nodes and Delaunay neighbors defined the edges. There was no distance threshold to the edge definition to avoid sub-graphs, long-distance edges were rare. For the baseline GCN model, the protein expression of each cell defines the node attributes (Fig. 3A). In the shape-aware GCN model, we concatenate the shape features vector to the cell type attributes. To perform a whole graph classification task we aggregated the information across nodes after the final layer, using global max pooling, to a single value of the clinical outcome. We used leave-one-out cross validation for TNBC, and 5-fold cross validation for HNC. The area under the Receiver Operating Characteristic (ROC) curve was used to compare the performance of models without versus with the shape priors. For the grade label in the MIBI-TNBC cohort, the data was imbalanced, thus we used micro-averaged ROC (i.e., weighted-average). Area under the ROC curve (AUC) was used as the measure of discrimination. Note that all prediction scores across each leave-out-out model were pooled for this analysis. This analysis reports a lower bound because different models can produce different scores.

### Models’ optimization and evaluation

The DNN and the GCN models were optimized using the Adam optimizer.

In all equations below, is the k-dimensional ground truth vector for a given cell, is the k-dimensional prediction for a given cell, is the per-dimension mean, is the number of cells, is the classes, k is the dimension of the GT and prediction vectors. The measures can be calculated per patient, or across the entire dataset.

For regression models, we used the mean squared error (MSE) loss:

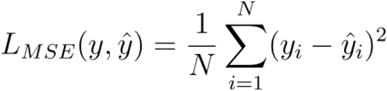

and performance was evaluated with the coefficient of determination R^2^:

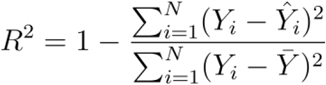

For classification models, we used the cross-entropy (CE) loss:

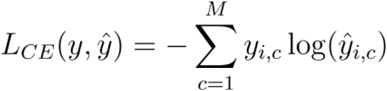

and performance was evaluated with the F1 score:

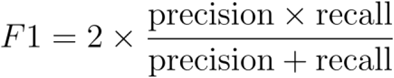

### Dimensionality reduction

We used PCA on the shape features for dimensionality reduction.

### Feature importance analysis (Explainability)

For AI explainability, we have used python’s Captum library, and specifically the GradientSHAP function. GradientSHAP is a gradient method to compute SHAP values, which are based on the Shapley values proposed in cooperative game theory and compute the contribution of each feature to each individual prediction (Lundberg & Lee, 2017). To cluster the feature importance patterns per protein, we used hierarchical clustering implementation in python’s SciPy package with the default settings.

## Statistical analysis

To test the significance of the contribution of the prior features to the target prediction, we performed a paired t-test comparing the R^2^ scores of patients between the baseline and the prior-aware model, to test the null hypothesis that the R^2^ differences are distributed around 0. One sample t-test was used in other cases.

## Supporting information

Appendix - Associating cell shape to cell type

Table S1 - data summary

Table S2 - IMC TNBC antibody panel

## Code and data availability

We are currently organizing our source code and processed data. We will make both publicly available as soon as possible (before journal publications). MIBI-melanoma and IMC-TNBC are part of larger cohorts that are currently still being collected and analyzed, and will be available upon request.

## Supplementary figures

**Figure S1.**
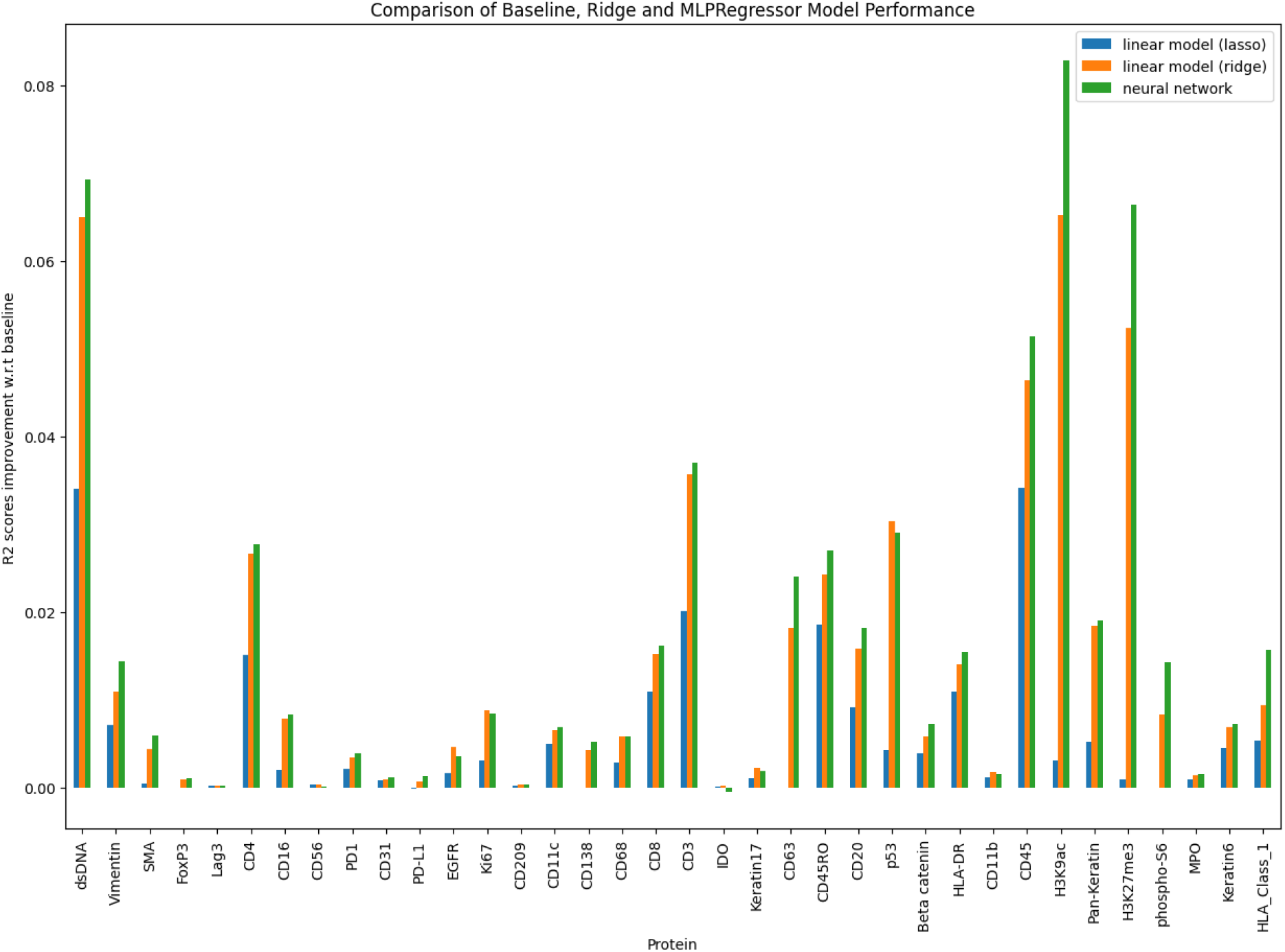
A shape-aware deep neural network (DNN) surpassed shape-aware linear models in the prediction of single cell protein expression. Comparison of the ΔR^2^ scores between a baseline model and three distinct shape-aware models—lasso (blue) and ridge (orange) linear models, and a fully connected DNN (green). The DNN consistently demonstrates superior performance over the linear models. Statistical significance for the improvement was calculated via paired t-tests comparing the R^2^ scores of proteins between the models to test the null hypothesis that the matched differences between the models are distributed around 0. p-value of 0.036 (DNN versus ridge) and p-value < 0.001(DNN versus lasso).

**Figure S2.**
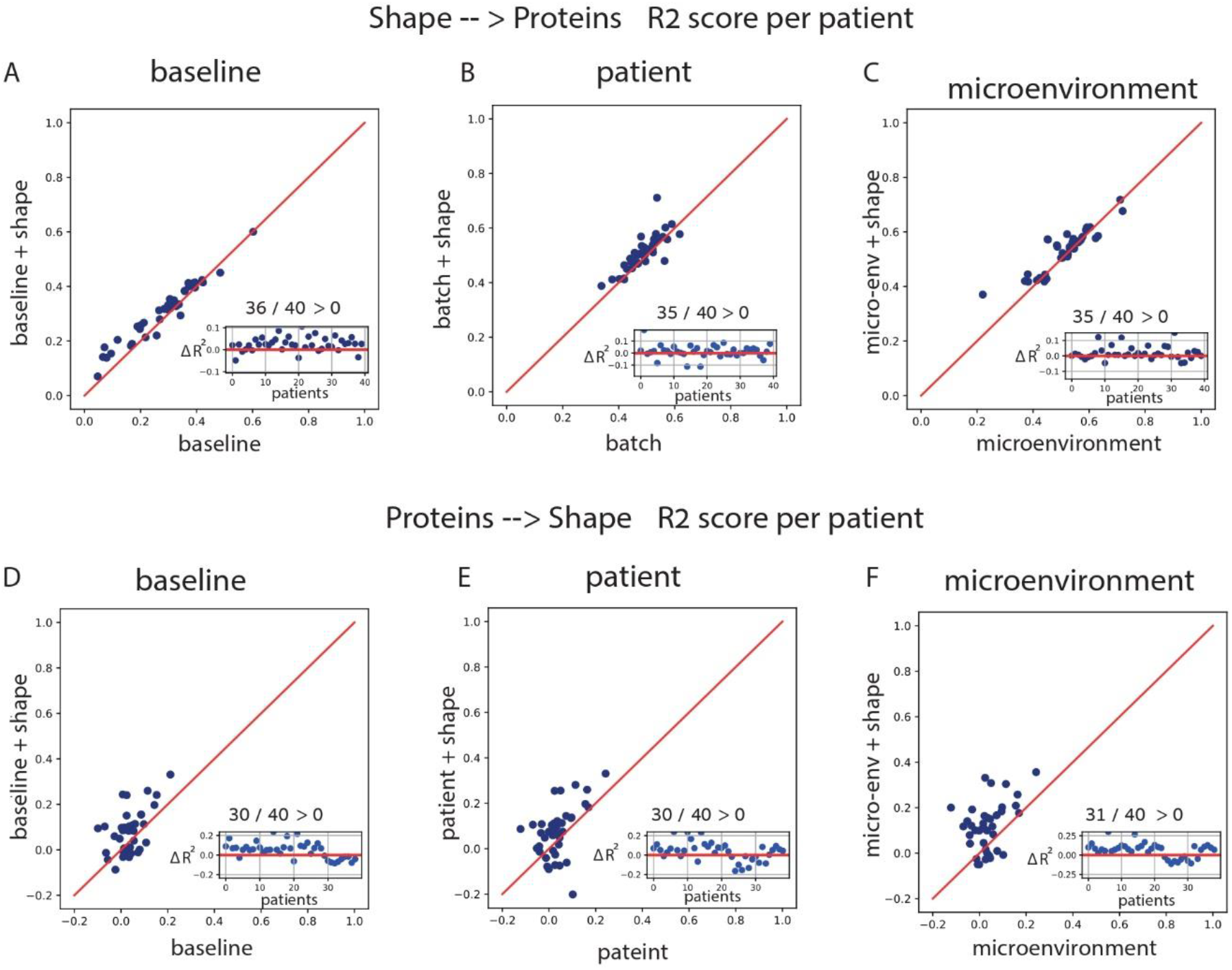
Each data point represents one patient’s R^2^ score, calculated based on the deviation of the predicted from the observed protein expression values, of a shape-aware (y-axis) and the baseline (x-axis) predictions. Data points above the diagonal y = x indicate more accurate shape-aware predictions. The inset shows the ΔR^2^ improvement by inclusion of shape per patient. Statistical significance is calculated via paired t-test, testing the null hypothesis that the R^2^ differences per patient are distributed around 0. **(A-C)** Shape-aware models improved the protein expression prediction accuracy. **(A)** Inclusion of shape improved a baseline that used the cell type for 36/40 of the patients, p-value = 0.027. **(B)** Inclusion of shape improved a baseline that used the cell type and the patient (batch term) for 35/40 of the patients, p-value = 0.034. **(C)** Inclusion of shape improved a baseline that used cell type, the patient and the cell’s microenvironment composition for 35/40 of the patients, p-value = 0.013. **(D-F)** Protein-aware models improved the shape prediction accuracy. **(D)** Inclusion of protein expression improved a baseline that used the cell type for 30/40 of the patients, p-value < 0.001. **(E)** Inclusion of protein expression improved a baseline that used the cell type and the patient (batch term) for 30/40 of the patients, p-value = 0.01. **(F)** Inclusion of protein expression improved a baseline that used the cell type, the patient and the cell’s microenvironment composition for 31/40 of the patients, p-value < 0.001.

**Figure S3.**
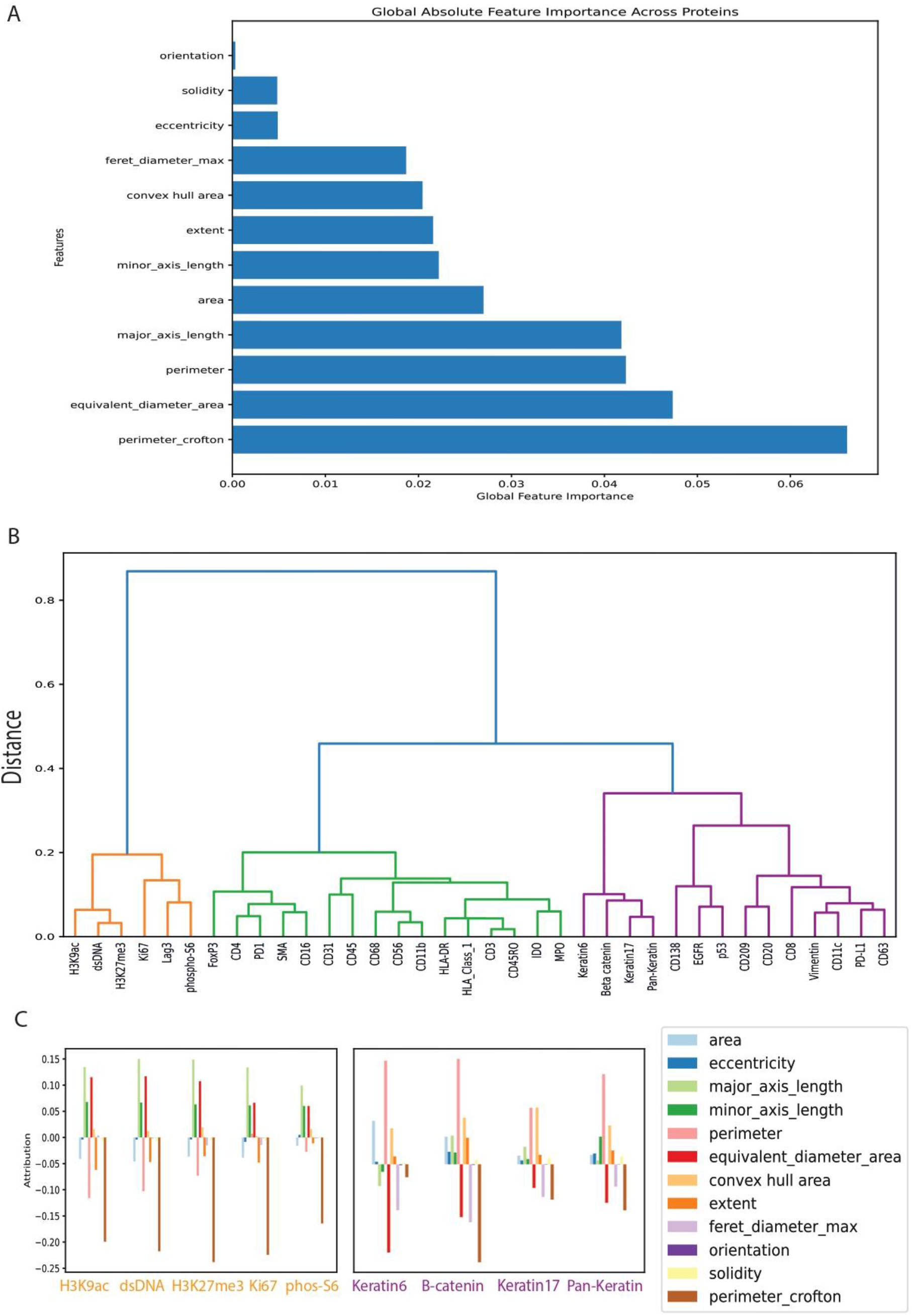
SHapley Additive exPlanations (SHAP) shape importance in protein prediction is clustered according to the proteins’ functions. **(A)** Global absolute shape importance accumulated across all cells showing the overall contribution of each shape feature to the protein expression prediction. Features are sorted by importance. **(B)** Hierarchical clustering of targets based on their feature importance profiles. **(C)** SHAP shapes importance for selected predicted proteins. Targets with similar functional roles exhibit similar patterns of feature importance.

**Figure S4.**
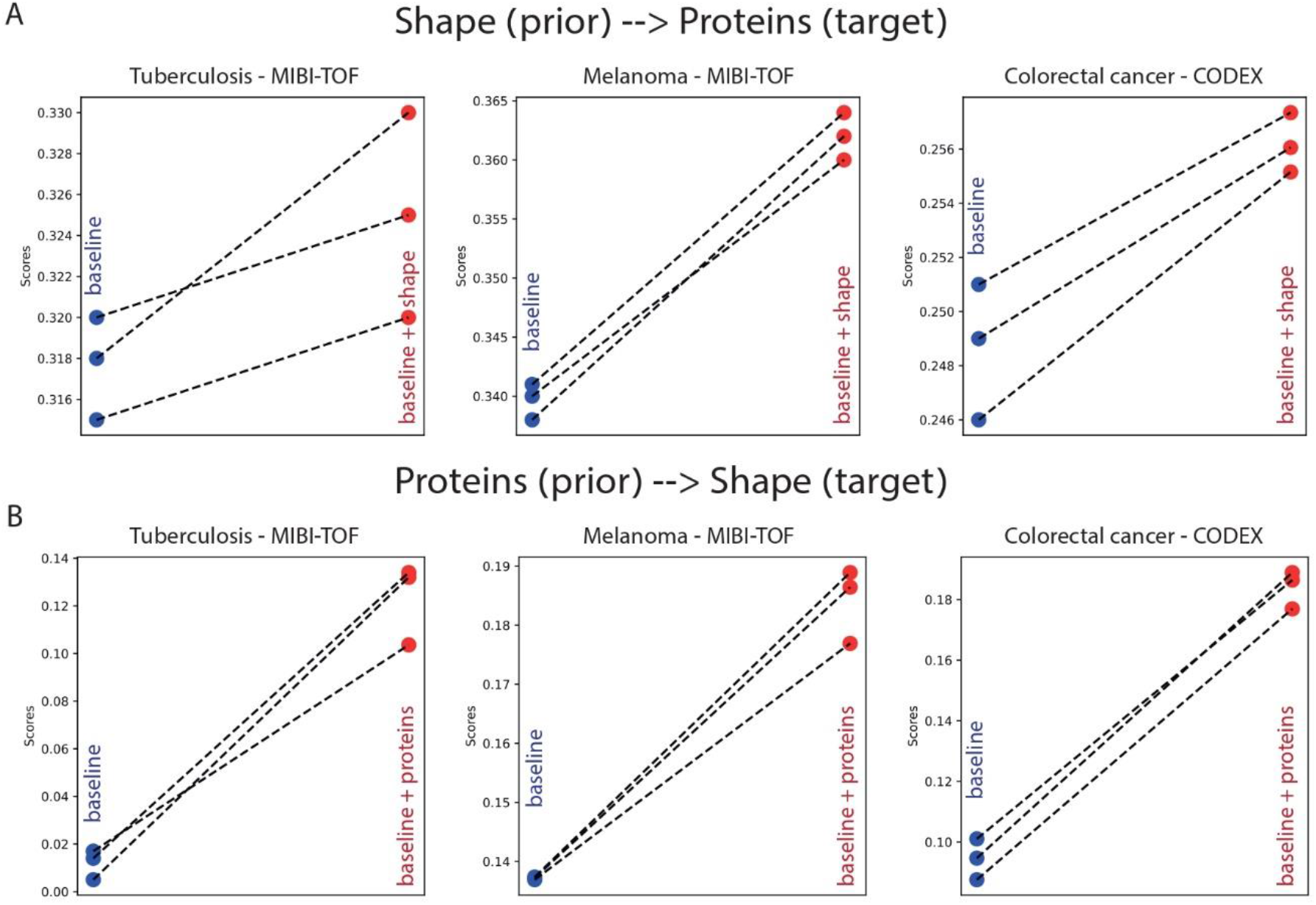
Shape- and protein-aware models surpassed their corresponding baseline models across diseases. Each matched data pair represents the 3-fold cross-validation R^2^ scores between the prediction and observed protein expression of the baseline (blue) and its matched shape-aware (A) or protein-aware (B) predictions (red). Statistical significance for the contribution of shape was calculated via paired t-tests comparing the R^2^ scores of patients between the baseline and the shape-aware model to test the null hypothesis that the matched differences between these models are distributed around 0. The null hypothesis was rejected with a p-value < 0.05 for all panels. Baseline models use the cell type versus the corresponding shape- and protein-aware models. Datasets (left-to-right): tuberculosis (MIBI-TOF), melanoma (MIBI-TOF), and colorectal cancer (CODEX).

**Figure S5.**
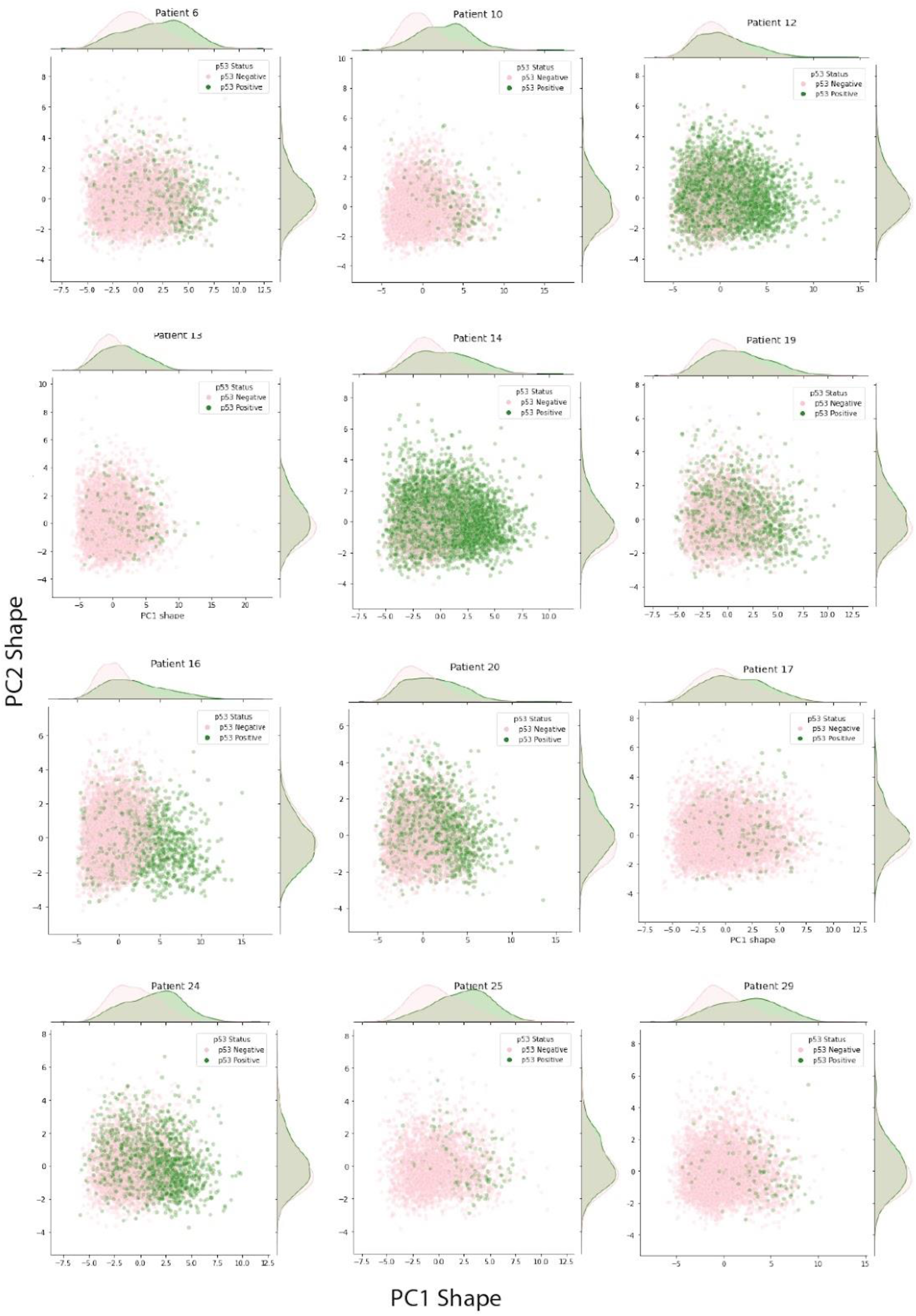
Shape differences between p53-positive and p53-negative tumor cells across multiple patients. **(A)** Scatter plots of tumor cells in the principal component analysis (PCA) shape-space, with cells colored based on their p53 status: positive (green) or negative (pink). The consistent separation between p53-positive and p53-negative cells in the PCA shape-space across multiple patients highlights the robustness of the association between p53 status and cell shape in tumor cells. T-test between PC1 distribution of the two groups is significant for all patients in this panel (p-value < 0.001).

**Figure S6.**
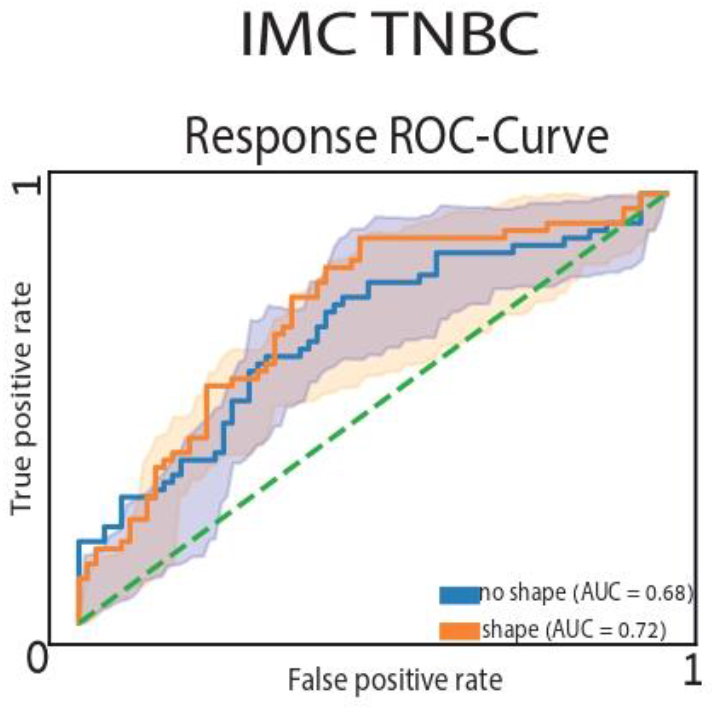
Shape contribution generalized to a different cohort. Mean (solid line) and standard deviation (shade) ROC-curve for a different TNBC dataset. The ROC curves compare the performance of shape-aware models (orange) and baseline models (blue) on 5-fold cross-validation. Inclusion of cell shape features led to an improvement of treatment response prediction.

## Funding and Acknowledgments

This research was supported by the Wellcome Leap Delta Tissue program (to AZ, LS, MP and CS).

## Author Contribution

AZ and LK conceived the study. YT developed the computational methods. YT, LK and AZ analyzed and interpreted the results. YT, OK and AZ drafted the manuscript. YB and CO generated data. LL and GT preprocessed data. AZ, LK, MP, LS and CS acquired funding and mentored YT, YB, CO, LL, GT, and LAR. All authors edited the manuscript and approved its content.

---

**Supplementary Table 1.** Datasets used in this study: MIBI-TNBC (L. Keren et al., 2018), MIBI-TB (McCaffrey et al., 2022), CODEX-HNC (Z. Wu et al., 2022), CODEX-CRC (Schürch et al., 2020), MIBI-Melanoma (Keren lab, Weizmann Institute), and IMC-TNBC (Parsons lab, King’s College London).

| Dataset | Technology | Disease | # patients | # cells | # proteins | # cell types | resolution |
| --- | --- | --- | --- | --- | --- | --- | --- |
| MIBI-TNBC | MIBI-TOF | TNBC | 40 | 178,366 | 36 | 16 | 0.5 $\mu\text{m}$ px |
| MIBI-TB | MIBI-TOF | TB | 26 | 56,402 | 38 | 19 | 0.5 $\mu\text{m}$ px |
| CODEX-HNC | CODEX | HNC | 139 | 2,061,162 | 22 | 16 | 0.4 $\mu\text{m}$ px |
| CODEX-CRC | CODEX | CRC | 35 | 218,372 | 56 | 16 | 0.4 $\mu\text{m}$ px |
| MIBI-Melanoma | MIBI-TOF | Melanoma | 69 | 1,701,560 | 36 | 27 | 0.5 $\mu\text{m}$ px |
| IMC-TNBC | IMC | TNBC | 58 | 3,738,846 | 35 | 32 | 1 $\mu\text{m}$ px |

