## Appendix - Associating cell shape to cell type for "Data-modeling the interplay between single cell shape, single cell protein expression, and tissue state"

### Cell shape variability in triple negative breast cancer (TNBC) tissues

We analyzed shape features heterogeneity from an existing MIBI-TOF human TNBC cohort, where every cell was segmented, assigned and expert-validated to one of 16 different cell types according to the cell's protein expression (Keren et al., 2018). We extracted 12 commonly used shape features from each cell (Appendix Table 1). Using the single cell type annotations, we analyzed shape features across the different cell types. Distributions of cell shape features pooled across patients according to their cell type showed large overlapping shape characteristics, identifying tumor cells as the largest (Appendix Fig. 1B). To overcome the confounding effect of inter-patient variability we compared the shape features distributions within each patient. First, for every cell type and for every shape feature we calculated the mean shape feature value per patient. Second, for each feature f and for each pair of cell types (X,Y), we tested the null hypothesis that the mean value of feature f in cell type X is the same as the mean value of feature f in cell type Y, using matched pairs of observations from the same patient. In other words, the null hypothesis states that the probability of an observation from cell type X being greater than an observation from cell type Y (P(Xfi > Yfi)) is equal to the probability of an observation from cell type Y being greater than an observation from cell type X (P(Yfi > Xfi)), where i denotes the patient from which the matched observations (i.e., mean value of feature f) are drawn. We applied the Wilcoxon test with FDR corrections for every triplet consisting of two cell types and a feature (*X*, *Y*, *f*), and examined pairs consisting of a cell type and a feature that were the largest / smallest among the other cell types. This analysis provided statistical evidence that, for the majority of patients, tumor cells exhibited the largest size, while macrophages demonstrated the smallest solidity (i.e, the ratio of the cell's area to its convex hull area) (Appendix Fig. 1C). However, most cell types did not have characteristic shape features that could distinguish them from the other cell types across patients implying that the association between cell type and cell shape was not straightforward.

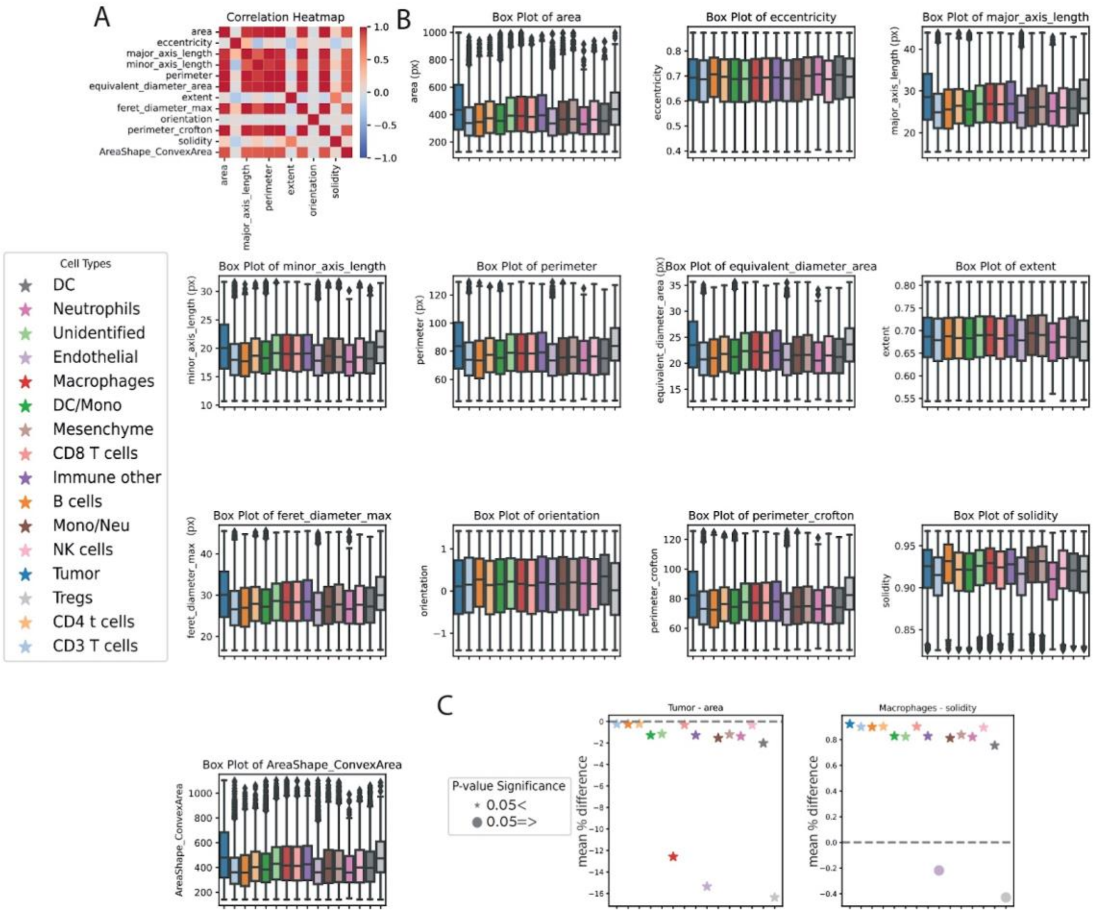


**Appendix Figure 1: Cell shape variability in triple negative breast cancer (TNBC) tissues**. (**A**) Correlation heatmap of 12 commonly used shape features extracted from each cell in the MIBI-TOF human TNBC cohort. The heatmap shows the extent of correlation between different shape features. (**B**) Box plots of various shape features (area, eccentricity, major axis length, minor axis length, perimeter, equivalent diameter area, extent, ferret diameter max, orientation, perimeter Crofton, solidity and convex area) for different cell types identified in the TNBC cohort. The distributions of cell shape features pooled across patients show large overlapping shape characteristics, with tumor cells being the largest. (**C**) Pairwise comparison of cell types for shape features within each patient using the Wilcoxon test with FDR corrections. The legend (left) shows the statistical significance (P-value) of the difference in the mean shape feature value between cell type pairs. Tumor cells exhibit the largest area, while macrophages demonstrate the smallest solidity across the majority of patients (right). However, most cell types do not have characteristic shape features that could distinguish them from other cell types across patients, suggesting that the association between cell type and cell shape is not straightforward.

### Cell shape contributes to cell type prediction at a low cell type hierarchies

To systematically and quantitatively explore more complex associations between cell type and cell shape we turned to using machine learning as a relative measurement metric. We used the single cell shape features to train XGBoost machine learning classifier (Chen & Guestrin, 2016) (from here forward referred to as 'model') to classify cells according to their types. The model was trained to predict cell-type from its shape features with a 80-20% split. The F1 score for each cell type ranged between 0-0.3, with a weighted average F1 score of 0.15, indicating that cell shape does not contain discriminative cell type information (Appendix Fig. 2A), at least at the crude spatial resolution of 0.5 $\mu$m microns per pixel of our data. To test the possibility that the weak link between cell type and cell shape was due to over specificity in classifying different cell types we performed the same machine learning analysis for an intermediate cell-lineage hierarchy that included four cell clusters: tumor, "immune adaptive", "immune non-adaptive", and "non-immune" (Appendix Fig. 2B). The intuition underlying this analysis was that the cell types may exhibit more pronounced differences in their shapes at this lower level of the cellular hierarchy. Even in the lower resolution of cell-lineage hierarchy, shape features showed weakly discriminating F1 scores ranging between 0.07-0.4, with a weighted average F1 score of 0.35 (Appendix Fig. 2C). These results indicate that cell shape alone does not contain strong discriminative information regarding the cell type, at least when determined using images acquired at the low spatial resolutions that are used in standard spatial single cell assays.

We speculated that although the high variability in cell shape confounded cell type prediction, the within cell type shape variability can contribute to protein expression-based cell type prediction. To test this hypothesis we trained XGBoost models to predict a cell's type from its protein expression, with or without inclusion of the cell's shape features. Shape-aware models deteriorated cell type classifications at the full spectrum of all 16 cell types (Appendix Fig. 2D), but improved cell type classification for the lower cell-lineage hierarchy of four cell clusters (Appendix Fig. 2E). These results suggest that (low resolution) cell shape can be associated to cell type only at lower lineage hierarchies.

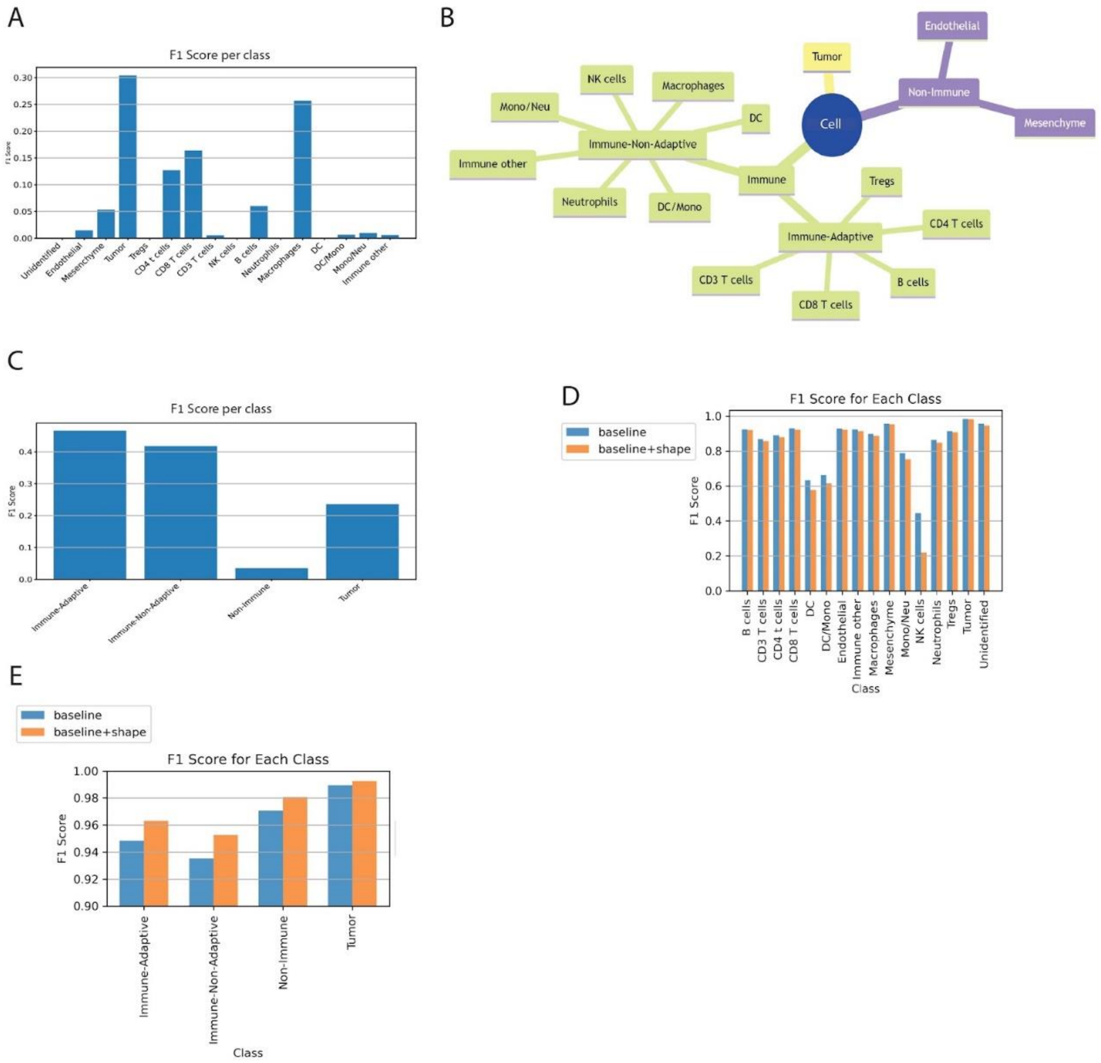


**Appendix Figure 2: Machine learning analysis of cell type and cell shape associations.** (**A**) F1 scores for predicting cell type from shape features using an XGBoost classifier. The model was trained on 80% of the data and tested on the remaining 20%. The F1 scores for each cell type range between 0-0.3, with a weighted average F1 score of 0.15, indicating that cell shape alone does not contain discriminative cell type information at the spatial resolution of 0.5 µm per pixel. (**B**) Schematic representation of the cell-lineage hierarchy used in the analysis, which includes four cell clusters: tumor, immune adaptive, immune non-adaptive, and non-immune. (**C**) F1 scores for predicting cell-lineage from shape features using the XGBoost classifier. Even at the lower resolution of cell-lineage hierarchy, shape features show weakly discriminating F1 scores ranging between 0.07-0.4, with a weighted average F1 score of 0.35. (**D**) Comparison of F1 scores for predicting cell type from protein expression features with (baseline+shape) and without (baseline) the inclusion of cell shape features. Shape-aware models deteriorate cell type classifications at the full spectrum of all 16 cell types. (**E**) Comparison of F1 scores for predicting cell-lineage from protein expression features with (baseline+shape) and without (baseline) the inclusion of cell shape features. Shape-aware models improve cell type classification for the lower cell-lineage hierarchy of four cell clusters, suggesting that cell shape can be associated with cell type only at lower lineage hierarchies when using low-resolution images.

### Cell area is most heterogeneous for tumor cells and most homogeneous for T-regulatory cells

While single cell shape did not hold much discriminative information regarding the cell type, we asked whether different cell types were characterized by different distributions of cell shape. We measured the coefficient of variation (CoV) in the cell area as a measure for heterogeneity. The CoV is defined as the ratio between an observable standard deviation to its mean in a given population, where larger CoV values indicate higher variability. We visualized the cell area CoV for each cell type and for each patient (Appendix Fig. 3A) and found that tumor cells' were the most heterogeneous and T-regulatory (‘Tregs’) cells’ were the most homogeneous cell types (Appendix Fig. 3B).

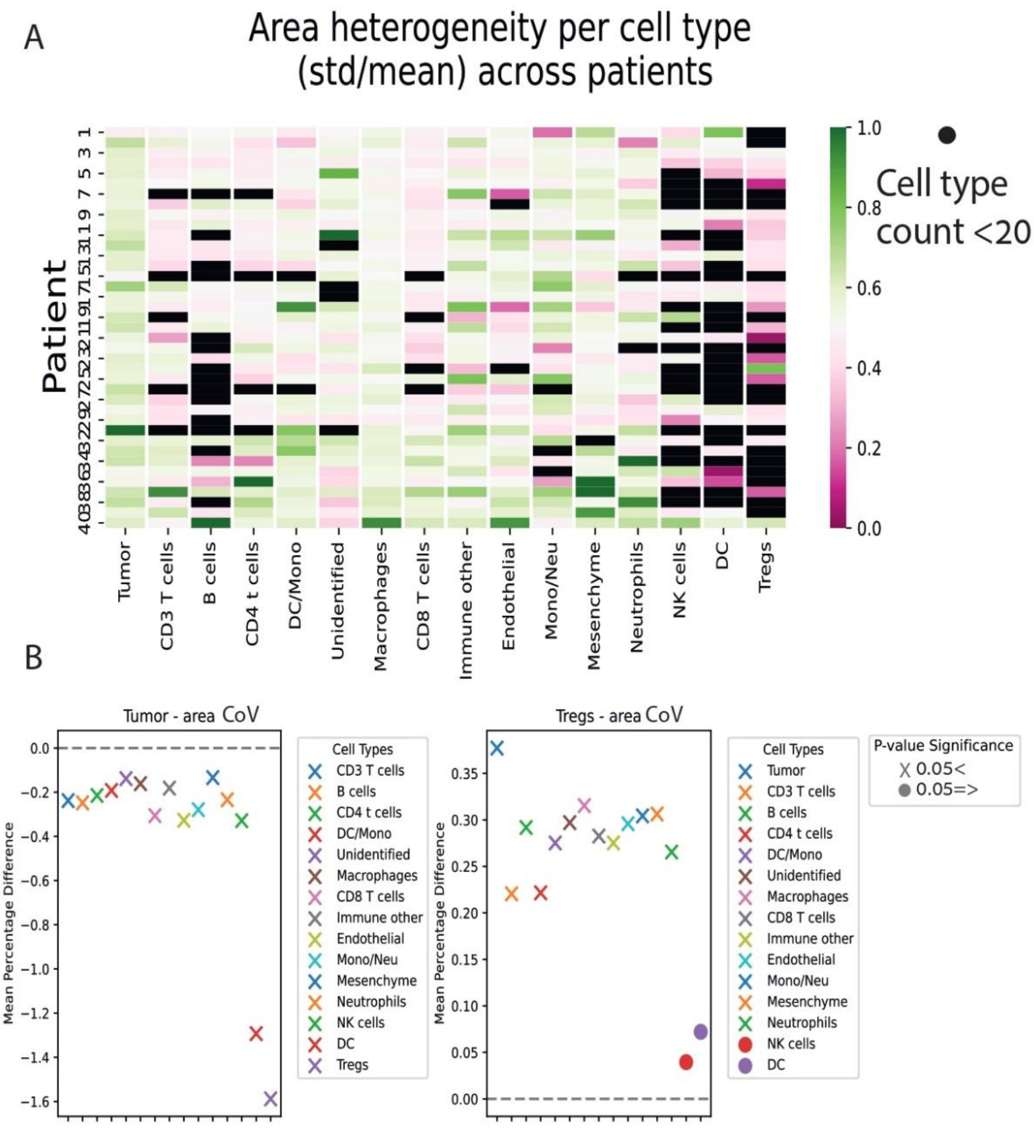

**Appendix Figure 3: Area heterogeneity per cell type across patients**. (**A**) Heatmap showing the coefficient of variation (CoV) of cell area for each cell type (columns) and patient (rows). The color scale represents the CoV values, with darker colors indicating higher heterogeneity. Cell types with a count less than 20 cells per patient are shown in black. (B) Pairwise comparison of cell types for area CoV within each patient using the Wilcoxon test. Tumor cells exhibit the highest heterogeneity in cell area, while Tregs cells show the lowest heterogeneity. The significance of the difference between tumor and Tregs to other cells is indicated by the P-value (Wilcoxon test), with $P < 0.05$ considered significant.

| Feature | Description |
|---|---|
| Area | The area (in μm$^2$) of the segmented cell mask |
| Eccentricity | The ratio of the focal distance over the major axis of the equivalent (i.e., that has the same normalized second central moments as the segmented cell mask) ellipse. The value is in the interval [0, 1), where a circle has an eccentricity of 0. |
| Major Axis Length | The length of the major axis of the ellipse that has the same normalized second central moments as the segmented cell mask. |
| Minor Axis Length | The length of the minor axis of the ellipse that has the same normalized second central moments as the segmented cell mask. |
| Perimeter | The curve length (in μm) around the shape's contour |
| Equivalent Diameter Area | The diameter of a circle with the same area as the segmented cell mask |

| | |
|---|---|
| Convex Hull Area | The area of the smallest convex polygon that encloses the segmented cell mask |
| Extent | The ratio between the area of the segmented cell mask and the area of its bounding box |
| Diameter Max | The largest distance between any two points in the segmented cell mask |
| Orientation | The angle between the x-axis and the major axis of the equivalent ellipse |
| Perimeter Crofton | The perimeter estimate using Crofton's formula |
| Solidity | The ratio of pixels in the region to pixels of the convex hull image |

Appendix Table 1: Shape features
